# ProSPr: Democratized Implementation of Alphafold Protein Distance Prediction Network

**DOI:** 10.1101/830273

**Authors:** Wendy M Billings, Bryce Hedelius, Todd Millecam, David Wingate, Dennis Della Corte

**Affiliations:** Department of Physics and Astronomy, Brigham Young University, Utah; Department of Computer Science,, Brigham Young University, Utah

## Abstract

Deep neural networks have recently enabled spectacular progress in predicting protein structures, as demonstrated by DeepMind’s winning entry with Alphafold at the latest Critical Assessment, of Structure Prediction competition (CASP13). The best protein prediction pipeline leverages intermolecular distance predictions to assemble a final protein model, but this distance prediction network has not been published. Here, we make a trained implementation of this network available to the broader scientific community. We also benchmark its predictive power in the related task of contact prediction against the CASP13 contact prediction winner TripletRes. Access to ProSPr will enable other labs to build on best in class protein distance predictions and to engineer superior protein reconstruction methods.

## Introduction

Recently, a variety of powerful protein structure prediction methods, based on machine learning algorithms, have been reported.[1] Although direct prediction of structure from sequence has been attempted,[2] reproducible success is currently based on two-stage protocols.[3] The first stage is the training of a deep convolutional neural network (CNN) that predicts some macromolecular structure restraints like residue to residue distances, residue contacts, dihedral angles or secondary structure assignments.[4] In a second stage, these restraints are used to construct a folded three-dimensional structure of the target protein. In the recent Critical Assessment of Structure Prediction (CASP13) a two stage folding protocol developed by DeepMind outperformed all established academic groups and predicted 25 of 43 protein structures with highest quality.[5] Unfortunately, DeepMind has not expressed a plan to publish the source code of their Alphafold protocol.

## Results & Discussion

Here, we report the re-implementation of the first part of the Alphafold pipeline, an intramolecular distance prediction CNN, made freely available as source code (https://github.com/dellacortelab/prospr) and a Docker6 container (see Methods). The CNN is in agreement with architectural details revealed by DeepMind at the December 2018 CASP13 conference (http://predictioncenter.org/casp13/doc/presentations/) and recently presented at a symposium at Washington University (https://www.youtube.com/watch?v=uQ1uVbrIv-Q); however, certain design decisions and hyperparameters were not shared in sufficient detail and required re-engineering. A graphical abstract of the CNN is given in Figure 1.

**Figure 1:**
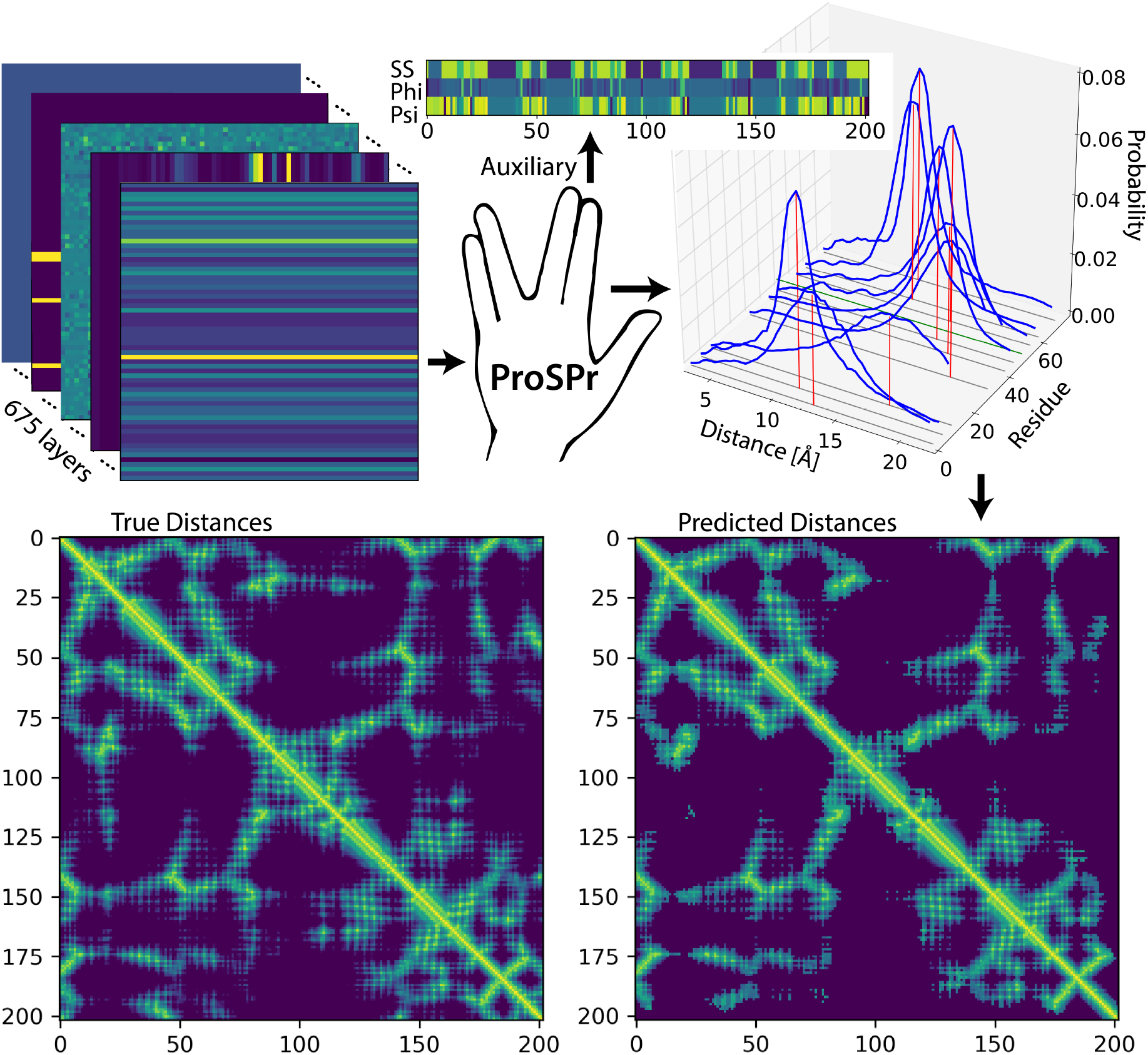
ProSPr distance prediction: Input sequence is converted into 675 layer input vector on top left. ProSPr CNN predicts as auxiliaries secondary structure elements for each residue from 9 DSSP classes (SS) and Phi/Psi Torsion angles between 0 and 360 degrees (top center). Further, it predicts distance distributions between each residue pair, shown as distance histogram for reside 50 of CASP Target T1016-D1 and selected residues (top right). The maxima of the distance distribution form a distance prediction map (heatmap, bottom right); left bottom, the real distances as measured in structure file of T1016-D1.

The CNN, named ProSPr (Protein Structure Prediction), predicts the *C_β_*–*C_β_* distance distributions between all amino acid residues (*C_α_* for Glycine) in a given protein sequence. We trained three versions of ProSPr on sequences in the CATH S35 dataset[7] (Supplementary Note and Figure S1) with the same network architecture but different input vectors. ProSPr follows Alphafold exactly and uses as input features the sequence information, the results of multiple sequence alignments (MSA) computed with PSI-BLAST[8] and HHblits,[9] as well as a Potts model[10, 11, 12] calculated from the MSA. ProSPr2 omits the Potts model, and ProSPr3 only uses the sequence information as input.

The performance of these three models was tested on the CASP 13 dataset for free and template-based models. The predicted distance distributions were converted into contact probabilities (distance between residues < 8 Å) and precision scores for three different classes of contacts were calculated according to the CASP assessment protocol.[13] ProSPr precision scores were directly compared to the performance of CASP 13 winning CNN TripletRes[14] and are shown in Figure 2 (Supplementary Table S1). Without being explicitly trained for this purpose, ProSPr predicts contacts for 109 tested CASP 13 domains with precision comparable to TripletRes over all classes, as shown in Table 1. Table 1 shows precision scores for ProSPr contacts with a maximum distance distribution < 8 Å, and for the full set of contacts independent of distribution maxima. For high confidence predictions, with maximum <8 Å, ProSPr is on average 2% better than Triplet Res on the L/5 scores. The L/2 and L scores are not directly comparable, because the absolute number of contacts ranked for ProSPr is substantially lower if the maximum < 8 Å criterion is applied than the total number of possible contacts ranked with TripletRes. For precision comparison the ranked probabilities of all contacts, independent of maximum, are therefore also reported. Under these conditions, we see that ProSPr is comparable to TripletRes. though on average slightly inferior. ProSPr2 results are comparable to ProSPr short and medium length contact predictions but are inferior to ProSPr long contact predictions. ProSPr3 is inferior to ProSPr in all categories. The performance of ProSPr2/3 was compared to TripletRes and is shown in Supplementary Figure S2. One issue with current precision reporting is that a smaller number of high confidence predictions leads to an inflation of L and L/2 scores, making model comparisons based on precision metric alone difficult to interpret. However, L/5 scores measure accurately the ability of a network to assign high confidence contacts and ProSPr outperforms TripleRes by an average of 2 %, which is in agreement with reports given by the Alphafold authors. (https://www.youtube.com/watch?v=uQ1uVbrIv-Q) Because ProSPr is trained to predict distances, the comparison against TripletRes only serves as a proof of concept. It would be a simple task to change the ProSPr network’s final layers and to train it explicitly for contact predictions, which was not the scope of this work.

**Figure 2:**
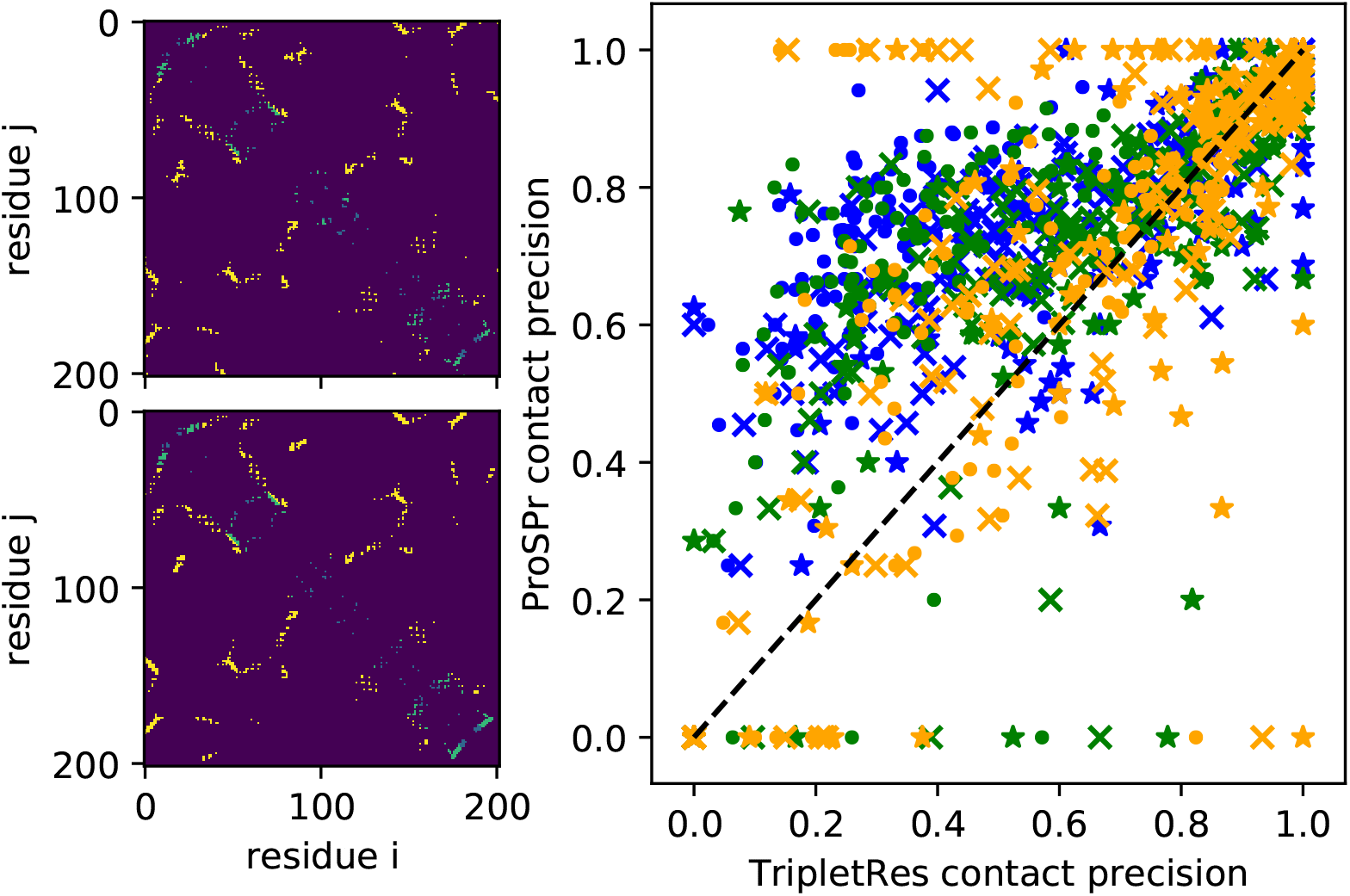
ProSPr distance predictions for 109 CASP13 domains were converted to contacts. Left panel shows an example contact label set for TR1016-D1 on top and the predicted contacts on the bottom. Right panel compares the precision for ProSPr contacts to those of TripletRes. Contacts are colored in blue, green, yellow for short, mid, and long. Markers circle, x, star correspond to L, L/2, L/5.

**Table 1:** Average precision scores for TripletRes and different ProSPr models are compared. ProSPr Average only ranks contacts where the maximum of the distance probability distribution falls between 0-8 Å, all other ProSPr rows sort contacts by total probability to be between 0-8 Å.

|  | Short(\|i - j\| > 5 & \|i-j\| < 12) |  |  | Medium(\|i - j\| > 11 & \|i-j\| < 24) |  |  | Long(\|i - j\| > 23) |  |  |
| --- | --- | --- | --- | --- | --- | --- | --- | --- | --- |
|  | L | L/2 | L/5 | L | L/2 | L/5 | L | L/2 | L/5 |
| TripletRes Average | 0.3001 | 0.4981 | 0.7276 | 0.3835 | 0.5787 | 0.7641 | 0.5308 | 0.6595 | 0.7627 |
| ProSPr Average | **0.6889** | **0.6938** | **0.7674** | **0.6680** | **0.6796** | **0.7657** | **0.6294** | **0.6972** | **0.7709** |
| ProSPr Full Average | 0.2980 | 0.4979 | 0.7428 | 0.3551 | 0.5511 | 0.7497 | 0.4969 | 0.6274 | 0.7436 |
| ProSPr2 Full Average | 0.2858 | 0.4639 | 0.6766 | 0.3378 | 0.4988 | 0.6746 | 0.3485 | 0.4459 | 0.5582 |
| ProSPr3 Full Average | 0.2064 | 0.2819 | 0.4018 | 0.2019 | 0.2587 | 0.3358 | 0.1287 | 0.1664 | 0.2176 |

Next to the python-based source code a Docker[6] container of ProSPr is made available to enable rapid usage of the distance prediction protocol. The container includes input vectors for select CASP13 targets, three pre-trained ProSPr models, and the distance prediction function to reproduce the results reported here. In addition, the distribution includes all dependencies necessary to produce a distance prediction for arbitrary sequences. Furthermore, the training set based on the CATH database, including the MSA and Potts models, is made available (https://byu.box.com/v/ProteinStructurePrediction) to repeat the training outlined in the methods section (approximately 2 TB of data). The GitHub repository contains a training function that can be used to either improve a pretrained model, or to train a modified ProSPr model for further optimization or ablation testing (full training on CATH dataset takes ~4 weeks on single T100 GPU). The original Alphafold protocol ensembled distance predictions over 4 separately trained models and subtracted a reference network during CASP13. A pretrained reference network is also provided that predicts distances only from sequence length and whether each residue is glycine (Supplementary Note). With time, we will make additional converged models of ProSPr and more comprehensive Docker containers available, to enable model ensembling.

The field of protein structure prediction has to tackle the challenge of protein reconstruction from geometric distance restraint distributions. During CASP 13 it became apparent that converting good distance predictions into chemically sound structures is still an unsolved problem. [4] ProSPr lowers the entrance barrier for academic labs and enables the community to quickly build on top of the internal coordinate predictions to develop improved protein reconstruction protocols. Further, we anticipate applications of ProSPr to investigate validity of evolutionary constraints as apparent from MSA. as ProSPr makes it possible to rapidly compare the effects of many single mutations on protein distances. These insights might also enable improved algorithms for in-silico drug discovery for mutated targets. In addition, we observed that ProSPr can interpolate distances between missing residues (Supplementary Figure S3), rendering it as a possible tool to support protein reconstruction from low resolution x-ray or cryo-EM data.[15]

In conclusion, we have demonstrated that ProSPr. a CNN based on the scarce details available for Alphafold, predicts residue-residue contacts with accuracy comparable to CASP13 winner TripletRes. ProSPr has the potential to propel protein structure prediction forward by democratizing the deep neural network and to empower directed evolution and protein reconstruction efforts.

## Methods

### Overview of ProSPr Architecture

Distance predictions within ProSPr can be made by calling distance prediction function, which consists of three steps as shown in Figure 3. Initially, a (L+32)x(L+32) profile is constructed for a sequence of length L using PSIBLAST, HHblits, a Potts model, and adding a frame of 32 bins as padding (Supplementary Note). Second, for a set 64×64 crops, defined by a stride parameter, of the profile an input vector with dimensions 675×64×64 is assembled. The input vector encodes the raw parameters, score, H parameters and Frobenius norm derived from the Potts model (total of 530 layers). Further, it contains two layers that hold the lists of residues for the crop, 42 layers for one-hot encoding of the sequence, 40 layers for a position specific substitution matrix (PSSM), 60 layers for the HHblits profile, and one layer for the sequence length. Third, the input layer is propagated through the CNN. After an initial batch norm, 1 dimensional convolution filters are applied to reshape the vector to a 128×64×64 matrix. This matrix is iterated 220 times through a residual network (RESNET) block that performs batch norming, applies the exponential linear unit (ELU) activation function, projects down to 64×64×64 dimensions, applies again batch norming and ELU, and then cycles through 4 different dilation filters. The dilation filters have sizes 1,2,4, and 8 and are applied with a padding of the same size to retain dimensionality. After a final batch norm, the matrix is projected up to 128×64×64 and an identity addition is performed. After 220 iterations the final matrix is subject to two 1 dimensional convolutions that reshape it into the final distance and auxiliary predictions. The auxiliaries predict 8 classes of secondary structure as defined within the DSSP classifications, and the phi and psi dihedrals for each residue; the angles are binned with 10 degrees resolution between 0 and 360. Due to possible gaps in the sequence, an additional classification bin is introduced for each auxiliary prediction that represents unassignable information. The auxiliary predictions were only used for training but could yield additional insights in ProSPr applications.

**Figure 3:**
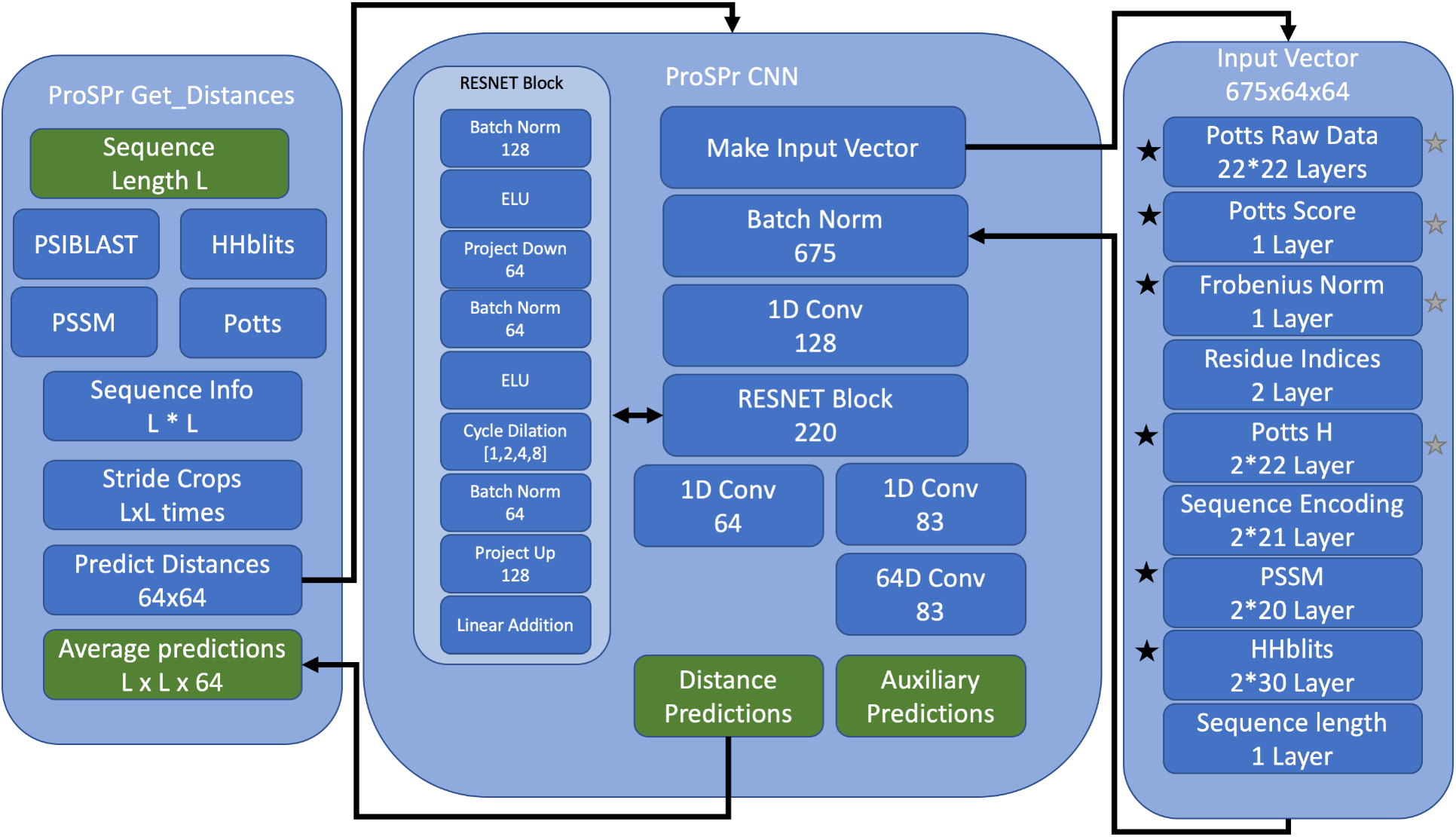
Overview of ProSPr core architectural components. On left the Get_Distances function, with inputs and outputs highlighted in green. In the center the ProSPr deep convolutional neural network with 220 RESNET blocks shown as an inlet. On the right a breakdown of input features, which were all used for ProSPr, the layers marked with a grey or black star were excluded in ProSPr2-3, respectively.

### Training of ProSPr

ProSPr was trained on 64×64 crops extracted from the CATH S35 dataset[7] with 26393, 1000, and 500 domains randomly selected as training, validation, and test sets, respectively (Supplementary Note). Initial weights were assigned randomly with Pytorch, the loss was calculated using cross entropy and an Adam optimizer with learning rate of 0.001 was used to update the weights. Total loss was calculated as the weighted sum of ten times the distance loss, the losses of two secondary structure assignme‘nts, and the losses of 4 torsion angles assignments. Training loss and validation loss converged after 500,000 iterations with training batch sizes of 8 (Supplementary Figure S1), which corresponds approximately to the number of total crops necessary to visit each subdomain in the training set once. The training of ProSPr2 and ProSPr3 used the same setup, only the input vectors contained different amount of information. For ProSPr2 all layers that contained Potts information were set to zero. For ProSPr3 the PSSM and HHBlits layers were also set to zero. For these networks, the training loss did not converge within 500,000 iterations (Supplementary Figure S1).

### Convert distances into contacts

As a test, the distances for 109 CASP13 domains, which were not included in the training or validation sets, were predicted and converted into contacts. Instead of using all possible 64×64 crops, a stride of 25 was used between the crops to speed up evaluation of large domains. Average contact scores improved by 1% when a stride of 1 was used for the 44 shortest domains. The 64×64×64 distance output encodes the probability of a residue i and j to have distances either not assignable (e.g. gap in sequence), in the range of 2.3 22 Å with. 3 Å resolution between classes, or greater than 22 Å. If the maximum of the probability distribution fell between 2.3 8 Å (bins 1-19), we considered two residues in contact for the high confidence predictions. In all cases, contacts were ranked according to the sum probability of distances between 2.3 and 8 Å and the top L, L/2, L/5 (L is length of sequence) contacts were selected to calculate accuracy scores. The contacts were classified based on the sequence separation of residues i and j into: short-range (6≤|i – j|≤11), medium-range (12≤|i – j|≤23) and long-range (|i – j|≥24) contacts.

### Evaluation of Contact Accuracy

According to CASP protocol, precision was calculated as follows:

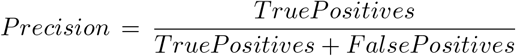

The average in each category was calculated over 109 test domains from CASP13. For the comparison with TripletRes, the difference in average precision per category was again averaged.

### Installation instruction for Docker

To install ProSPr as a docker container and to see all currently available options enter in the command line (after installing docker):

> docker run prospr/prospr

Yes, it is that easy!

## Acknowledgements

DDC expresses gratitude for computational resources offered by BYU Office of Research Computing.

## Supplementary Note

### Dataset Construction and Cleaning

Protoin domains and structures used in training ProSPr were obtained from the CATH s35 sequence homology dataset. A list of all domains in the s35 cluster with their sequences in FASTA format was downloaded from ftp://orengoftp.biochem.ucl.ac.uk/cath/releases/latest-release/cath-classification-data/cath-domain-list-S35.txt on 16 February 2019: domain names were then extracted from the text and their corresponding structural files downloaded individually using the following link: http://www.cathdb.info/vorsion/v4_1_0/api/rest/id/DOMAIN_ID.pdb. Some of the domains were not successfully downloaded, and brief manual inspection showed that structural files did not seem to exist for at least several of the domains specified in the s35 sequence list. Domains for which a structural file could not be obtained for any reason were excluded from the dataset.

Amino acid sequences for each domain were derived from the corresponding structure file. Although each domain was originally extracted from a FASTA file including the sequences, discrepancies existed between those explicit sequences and the string of residues contained in each structure file (e.g. some structures contained only portion(s) of the sequences specified separately). Additionally, monitoring of residue numbers present in the structure files enabled us to denote gaps in the sequences with a unique character, whereas no such notation existed in the original FASTA sequences.

For reasons further contextualized under Crops and Input Padding, we trimmed the beginning(s) and/or end(s) of protein sequences if gaps larger than or equal to 32 residues separated small terminal segments of fewer than 17 residues from the remainder of the protein. This eliminated instances in which possibly hundreds of residues whose identities and/or positions were undetermined in the protein structure (gaps) were included in the structure-derived sequence because several adjacent terminal residues are recorded (see CATH domain 1hu3A00 as example). In such cases, the characters corresponding to these end residues and the adjacent gap section(s) were removed from sequence, and the process was repeated for the remaining sequence until no additional changes were made. At the conclusion of these sequence modifications, corresponding structure files were trimmed to match the shorter sequences.

PSSMs were constructed using PSIBLAST (ftp://ftp.ncbi.nlm.nih.gov/blast/executables/blast+/LATEST/) and the nonredundant (nr) database, while alignments were generated using HHBlits from HHSuite (https://github.com/soedinglab/hh-suite) in conjunction with the uniclust30_2018_08 database (http://www.user.gwdg.de/~compbiol/uniclust/2018_08/.UNICLUST). Both ran with E-values of 0.001 and completed 3 iterations. A limit of 100000 was imposed on the number of sequences HHBlits could write out to the alignment file. HHBlits alignment results were then processed further and used in computing the Potts models (original obtained from https://github.com/magnusekeberg/plmDCA. however modifications were made to the code and those modifications are in our github repository for this project under src/potts.patch). Further information about these programs, as well as details concerning how the values of interest were extracted from these three sets of output files and used in the input vectors can be found in the code documentation.

Training labels for the distance prediction task as well as the auxiliary secondary structure and torsion angle predictions were created by making structural calculations (in the case of the auxiliary predictions, aided by the DSSP algorithm, code available at https://github.com/cmbi/xssp) and binning the observations, thus enabling ProSPr to treat each as a classification task. Pairwise distances between all available beta carbon atoms (except for alpha carbons in the case of glycine) were classified into 64 classes, where 62 represented equivalently-sized bins over the 2-22Å range (~0.32Å width each), one represented all distances greater than 22Å, and another signified a gap or missing part of the protein structure. Secondary structure classifications as made by DSSP were retained as separate classes, with the addition of one extra bin to again represent missing data (eg. no residuo(s) in that part of the structure) for a total of 9 classes. Phi and psi torsion angle values were classified into 36 10° bins ranging from −180° to 180° and one for gaps, resulting in a total of 37 possible classes.

After all inputs and labels were generated for available domains, an intersection was performed to extract the list of domains for which each of these independent tasks had executed successfully; this list was subsequently divided into the training, validation, and test sets as described in Methods.

**Figure 4:**
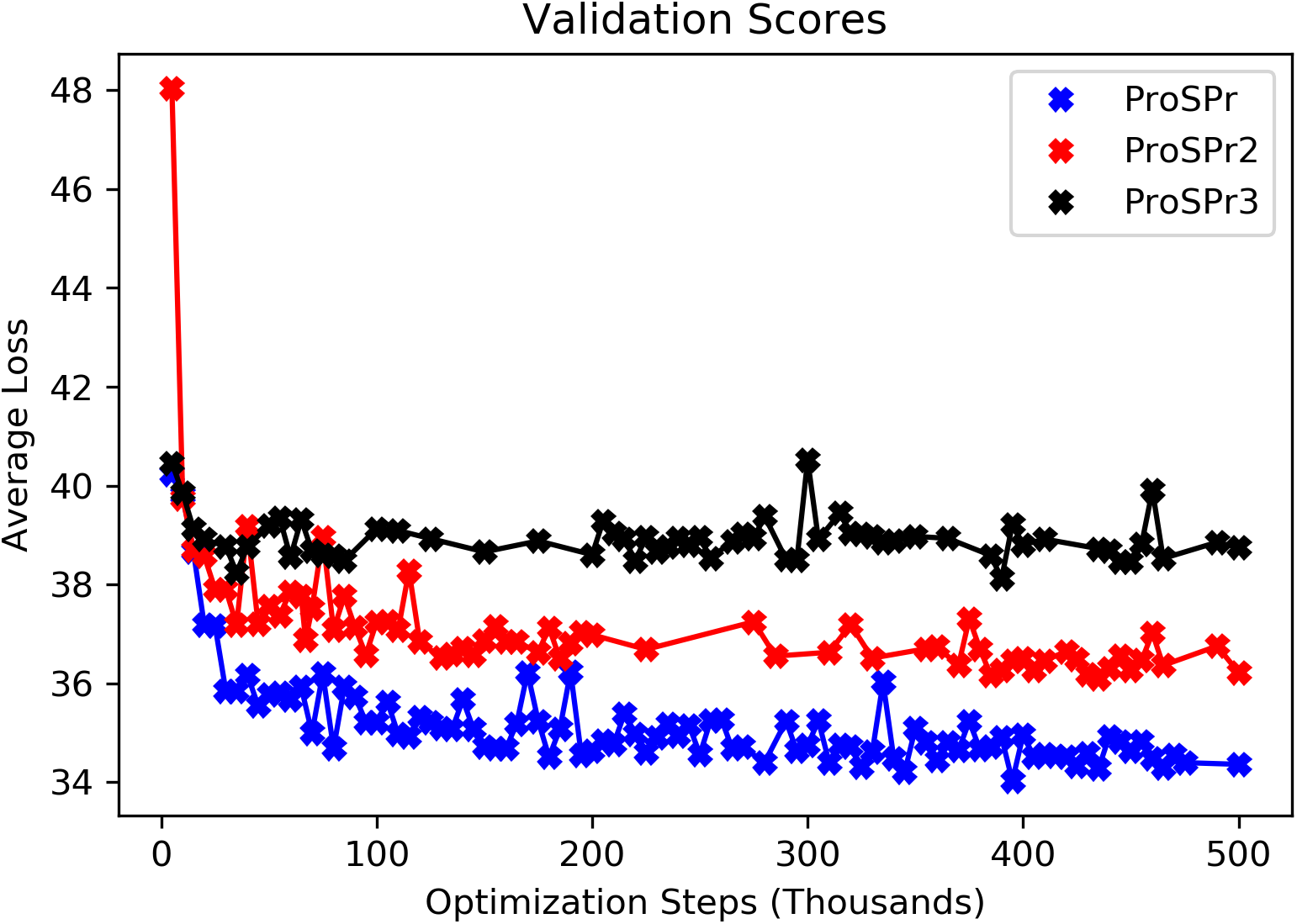
Supplementary Figure S1 Validation curves for ProSPr networks. For a batch size of 8 500,000 training iterations were conducted for full input vectors (ProSPr). input vectors without Potts Models (ProSPr2), and input vectors without any features derived from multiple sequence alignment (ProSPr3).

**Figure 5:**
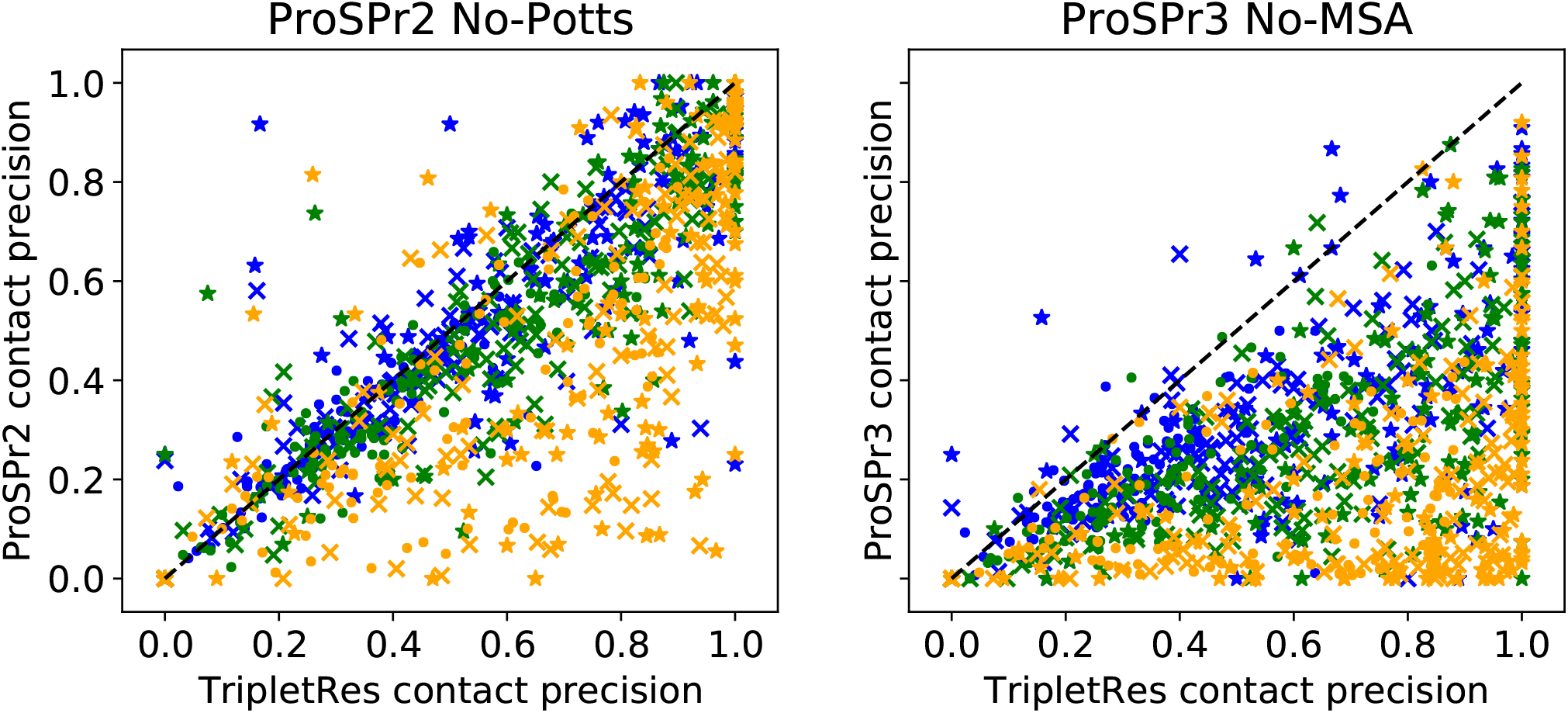
Supplementary Figure S2 Comparison of ProSPr2 and ProSPr3 contact precision against TripletRes for 109 CASP 13 domains. Contacts are colored in blue, green, yellow for short, mid, and long, respectively. Markers circle, x, star correspond to L, L/2, L/5.

**Figure 6:**
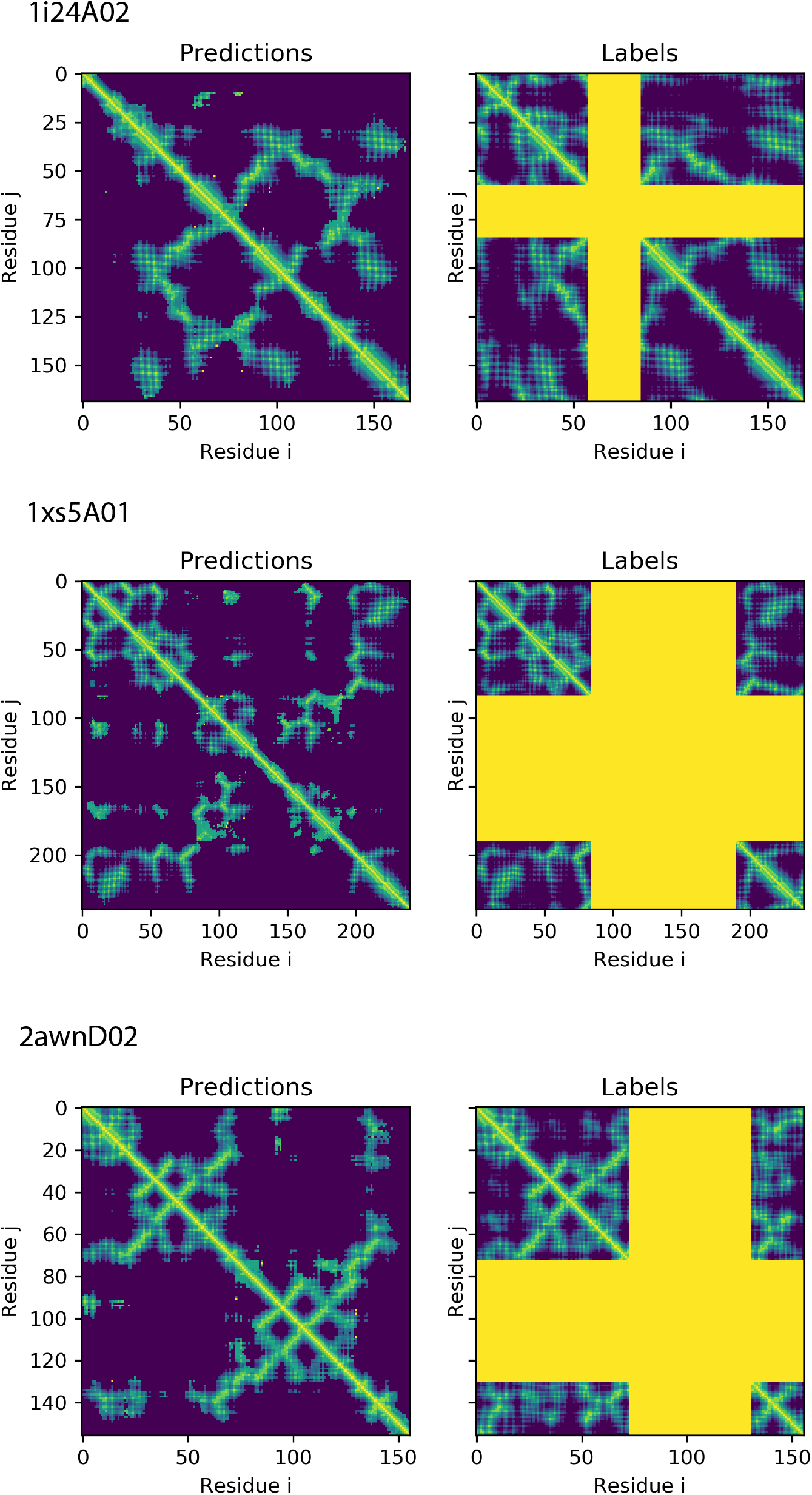
Supplementary Figure S3 ProSPr interpolates distance information for missing residues. For 3 sequences with missing residues from the CATH validation set we observed that ProSPr predicts distances for missing residues (yellow crosses in label). This might be a useful feature to enhance structure reconstruction efforts and is subject to further analysis.

### Crops and Padding

Using the dataloader function included and documented in the project source code, training is performed on 64×64 amino acid crops of training domains, specified using i and j (coordinate) options and the protein domain id. The i and j coordinates correspond to the amino acid positions on which the crop is centered, and can range from 0 to one less than the sequence length of that domain. This results in padding of up to 32 being added in either dimension so that the crop maintains its 64×64 size. Training on crops allows the network to use more training data than if the entire domains were used, and the consistent size helps in distributed training.

However, in testing and application of such a network, distance predictions for the entire protein domain are often of much greater relevance than those made for a single 64×64 residue crop. Therefore, outside of training (including for validation set testing, the analysis done with the CASP13 targets, etc.) full-domain predictions are made by processing multiple adjacent crops of the same domain and averaging those values where the crops overlap. The “stride” parameter specifies how far apart the i,j, coordinates for each crop are (eg. stride of 1 means that every possible i,j combination for the length of the protein will be processed), however the evaluation time increases as the square of the protein length. On the contacts data set we observed only small improvements of *1%* by reducing the stride from 25 to 1.

**Supplementary Table S1.** ProSPr contact scores for 109 CASP13 targets.

|  | Target | Short L | Short L/2 | Short L/5 | Mid L | Mid L/2 | Mid L/5 | Long L | Long L/2 | Long L/5 |
| --- | --- | --- | --- | --- | --- | --- | --- | --- | --- | --- |
| 0 | T0949-D1 | 0.5652 | 0.5652 | 0.5652 | 0.7727 | 0.7727 | 1.0000 | 0.8841 | 0.9565 | 1.0000 |
| 1 | T0950-D1 | 0.2500 | 0.2500 | 0.2500 | 0.5862 | 0.5862 | 0.5862 | 0.3879 | 0.3879 | 0.5441 |
| 2 | T0951-D1 | 0.7358 | 0.7358 | 0.7358 | 0.6667 | 0.6667 | 0.7358 | 0.7962 | 0.9167 | 0.9623 |
| 3 | T0953s1-D1 | 0.6667 | 0.6667 | 0.7692 | 0.7143 | 0.7143 | 0.7143 | 0.0000 | 0.0000 | 0.0000 |
| 4 | T0953s2-D1 | 0.6000 | 0.6000 | 0.6250 | 0.4615 | 0.4615 | 0.6250 | 1.0000 | 1.0000 | 1.0000 |
| 5 | T0953s2-D2 | 0.4000 | 0.4000 | 0.4000 | 0.0000 | 0.0000 | 0.0000 | 0.7500 | 0.8333 | 1.0000 |
| 6 | T0953s2-D3 | 0.8000 | 0.8000 | 0.8000 | 0.0000 | 0.0000 | 0.0000 | 0.0000 | 0.0000 | 0.0000 |
| 7 | T0954-D1 | 0.7614 | 0.8204 | 0.9394 | 0.6914 | 0.7904 | 0.9545 | 0.2680 | 0.3174 | 0.4394 |
| 8 | T0955-D1 | 0.6111 | 0.6111 | 0.7500 | 0.7500 | 0.7500 | 0.8750 | 0.0000 | 0.0000 | 0.0000 |
| 9 | T0957s1-D1 | 0.8000 | 0.8000 | 0.8571 | 0.8421 | 0.8421 | 0.9048 | 1.0000 | 1.0000 | 1.0000 |
| 10 | T0957s1-D2 | 0.5000 | 0.5000 | 0.5000 | 0.8000 | 0.8000 | 0.8000 | 0.9231 | 0.9231 | 1.0000 |
| 11 | T0957s2-D1 | 0.7619 | 0.7619 | 0.7619 | 0.7222 | 0.7222 | 0.7000 | 0.6842 | 0.6842 | 0.6842 |
| 12 | T0958-D1 | 0.3077 | 0.3077 | 0.3077 | 0.3636 | 0.3636 | 0.3333 | 0.4783 | 0.4783 | 0.4667 |
| 13 | T0959-D1 | 0.5000 | 0.5000 | 0.5429 | 0.5890 | 0.5890 | 0.6000 | 0.6963 | 0.7727 | 0.9429 |
| 14 | T0960-D1 | 0.0000 | 0.0000 | 0.0000 | 0.0000 | 0.0000 | 0.0000 | 0.0000 | 0.0000 | 0.0000 |
| 15 | T0960-D2 | 0.7333 | 0.7333 | 0.6875 | 0.7500 | 0.7500 | 0.9375 | 0.1667 | 0.1667 | 0.1667 |
| 16 | T0960-D3 | 0.8293 | 0.8293 | 0.8824 | 0.8488 | 0.9091 | 1.0000 | 0.6364 | 0.6364 | 0.7647 |
| 17 | T0960-D4 | 0.5000 | 0.5000 | 0.5833 | 0.2857 | 0.2857 | 0.2857 | 0.0000 | 0.0000 | 0.0000 |
| 18 | T0960-D5 | 0.6667 | 0.6667 | 0.7500 | 0.6538 | 0.6538 | 0.8000 | 0.8750 | 1.0000 | 1.0000 |
| 19 | T0961-D1 | 0.8348 | 0.8348 | 0.8600 | 0.7962 | 0.7962 | 0.9100 | 0.7789 | 0.9163 | 0.9600 |
| 20 | T0962-D1 | 0.8039 | 0.8039 | 0.8857 | 0.7333 | 0.7333 | 0.8000 | 0.6484 | 0.6932 | 0.7714 |
| 21 | T0963-D1 | 0.0000 | 0.0000 | 0.0000 | 0.0000 | 0.0000 | 0.0000 | 0.0000 | 0.0000 | 0.0000 |
| 22 | T0963-D2 | 0.7200 | 0.7200 | 0.6875 | 0.7407 | 0.7407 | 0.9375 | 1.0000 | 1.0000 | 1.0000 |
| 23 | T0963-D3 | 0.7857 | 0.7857 | 0.9444 | 0.8022 | 0.8478 | 1.0000 | 0.6786 | 0.6786 | 0.7222 |
| 24 | T0963-D4 | 0.5833 | 0.5833 | 0.5833 | 0.0000 | 0.0000 | 0.0000 | 0.0000 | 0.0000 | 0.0000 |
| 25 | T0963-D5 | 0.8095 | 0.8095 | 1.0000 | 0.6939 | 0.6957 | 0.8333 | 0.9247 | 1.0000 | 1.0000 |
| 26 | T0964-D1 | 0.7742 | 0.8298 | 0.9444 | 0.7959 | 0.8085 | 0.9444 | 0.8478 | 0.8936 | 1.0000 |
| 27 | T0965-D1 | 0.6667 | 0.6667 | 0.6774 | 0.6600 | 0.6600 | 0.8548 | 0.7981 | 0.8974 | 0.9355 |
| 28 | T0966-D1 | 0.6410 | 0.6410 | 0.6410 | 0.6620 | 0.6620 | 0.6620 | 0.2933 | 0.5429 | 0.8367 |
| 29 | T0967-D1 | 0.7200 | 0.7200 | 0.8667 | 0.8511 | 0.9231 | 1.0000 | 0.7949 | 0.8974 | 0.8000 |
| 30 | T0968s1-D1 | 0.7234 | 0.7234 | 0.9130 | 0.8750 | 0.8983 | 1.0000 | 0.4348 | 0.5254 | 0.8261 |
| 31 | T0968s2-D1 | 0.8256 | 0.9298 | 1.0000 | 0.7037 | 0.7544 | 0.9565 | 0.2500 | 0.2500 | 0.3043 |
| 32 | T0969-D1 | 0.6364 | 0.6364 | 0.6857 | 0.6914 | 0.6914 | 0.7286 | 0.6969 | 0.8693 | 1.0000 |
| 33 | T0970-D1 | 0.8125 | 0.8125 | 0.8125 | 0.8235 | 0.8235 | 0.8947 | 0.7742 | 0.7742 | 0.8947 |
| 34 | T0971-D1 | 0.8036 | 0.8036 | 0.9600 | 0.8679 | 0.9688 | 1.0000 | 0.7364 | 0.9531 | 1.0000 |
| 35 | T0973-D1 | 0.7500 | 0.7500 | 0.7500 | 0.8182 | 0.8182 | 0.8800 | 0.6176 | 0.6774 | 0.9200 |
| 36 | T0974s1-D1 | 0.7692 | 0.7692 | 1.0000 | 0.8333 | 0.8333 | 0.8333 | 0.7037 | 0.7037 | 0.9231 |
| 37 | T0974s2-D1 | 0.7879 | 0.7879 | 1.0000 | 0.7619 | 0.7619 | 0.8667 | 0.7593 | 0.7949 | 0.9333 |
| 38 | T0975-D1 | 0.7375 | 0.7375 | 0.7931 | 0.5824 | 0.5824 | 0.7759 | 0.4658 | 0.6507 | 0.7414 |
| 39 | T0976-D1 | 0.7600 | 0.7600 | 0.8261 | 0.8444 | 0.8444 | 1.0000 | 0.8824 | 0.9661 | 1.0000 |
| 40 | T0976-D2 | 0.7308 | 0.7308 | 0.7917 | 0.8750 | 0.8750 | 1.0000 | 0.8537 | 0.9344 | 1.0000 |
| 41 | T0977-D1 | 0.8417 | 0.8417 | 0.9167 | 0.8148 | 0.8933 | 0.9500 | 0.3898 | 0.3898 | 0.5333 |
| 42 | T0977-D2 | 0.7143 | 0.7143 | 0.7143 | 0.8511 | 0.8511 | 0.8500 | 0.7143 | 0.7143 | 0.7143 |
| 43 | T0978-D1 | 0.6667 | 0.6667 | 0.6667 | 0.6629 | 0.6629 | 0.7073 | 0.4272 | 0.6796 | 0.9146 |
| 44 | T0979-D1 | 0.0000 | 0.0000 | 0.0000 | 0.0000 | 0.0000 | 0.0000 | 0.0000 | 0.0000 | 0.0000 |
| 45 | T0980s1-D1 | 0.7500 | 0.7500 | 0.8500 | 0.5714 | 0.5714 | 0.5714 | 0.6000 | 0.6000 | 0.6000 |
| 46 | T0980s2-D1 | 0.6000 | 0.6000 | 0.6000 | 0.0000 | 0.0000 | 0.0000 | 0.0000 | 0.0000 | 0.0000 |
| 47 | T0981-D1 | 0.8571 | 0.8571 | 0.9412 | 0.8182 | 0.8095 | 0.8824 | 0.5000 | 0.5000 | 0.5000 |
| 48 | T0981-D2 | 0.6250 | 0.6250 | 0.6000 | 0.7143 | 0.7143 | 0.7143 | 1.0000 | 1.0000 | 1.0000 |
| 49 | T0981-D3 | 0.5500 | 0.5500 | 0.5500 | 0.7647 | 0.7647 | 0.7647 | 0.8168 | 0.9406 | 1.0000 |
| 50 | T0981-D4 | 0.8750 | 0.8750 | 0.8750 | 0.8824 | 0.8824 | 0.8824 | 1.0000 | 1.0000 | 1.0000 |
| 51 | T0981-D5 | 0.6522 | 0.6522 | 0.9200 | 0.7419 | 0.7419 | 0.7600 | 0.8034 | 0.9048 | 1.0000 |
| 52 | T0982-D1 | 0.8169 | 0.8507 | 1.0000 | 0.7879 | 0.8955 | 0.9615 | 0.7188 | 0.9104 | 1.0000 |
| 53 | T0982-D2 | 0.7302 | 0.7302 | 0.8846 | 0.6627 | 0.7538 | 0.9615 | 0.6552 | 0.7846 | 0.8077 |
| 54 | T0983-D1 | 0.8261 | 0.8261 | 0.8298 | 0.8099 | 0.8034 | 1.0000 | 0.7787 | 0.8974 | 0.9787 |
| 55 | T0984-D2 | 0.4545 | 0.4545 | 0.4545 | 0.3333 | 0.3333 | 0.3333 | 0.3774 | 0.3774 | 0.4828 |
| 56 | T0986s1-D1 | 0.7455 | 0.8222 | 1.0000 | 0.6923 | 0.6923 | 0.6667 | 0.6800 | 0.6800 | 0.8333 |
| 57 | T0986s2-D1 | 0.6531 | 0.6531 | 0.7333 | 0.8108 | 0.8108 | 0.8667 | 0.3226 | 0.3226 | 0.3333 |
| 58 | T0987-D1 | 0.8438 | 0.8438 | 0.8438 | 0.6279 | 0.6279 | 0.6389 | 0.7273 | 0.7273 | 0.8889 |
| 59 | T0987-D2 | 0.6053 | 0.6053 | 0.6053 | 0.6000 | 0.6000 | 0.6000 | 0.5972 | 0.5972 | 0.7073 |
| 60 | T0989-D1 | 0.5581 | 0.5581 | 0.5000 | 0.6032 | 0.6032 | 0.7308 | 1.0000 | 1.0000 | 1.0000 |
| 61 | T0989-D2 | 0.9412 | 0.9412 | 0.9412 | 0.8000 | 0.8000 | 0.8000 | 0.0000 | 0.0000 | 0.0000 |
| 62 | T0990-D1 | 0.6923 | 0.6923 | 0.6923 | 0.8000 | 0.8000 | 0.8000 | 0.5882 | 0.5882 | 0.7333 |
| 63 | T0990-D2 | 0.6512 | 0.6512 | 0.6512 | 0.6531 | 0.6531 | 0.6889 | 0.6071 | 0.6071 | 0.6071 |
| 64 | T0990-D3 | 0.4571 | 0.4571 | 0.4571 | 0.5313 | 0.5313 | 0.5313 | 0.6279 | 0.6279 | 0.6429 |
| 65 | T0992-D1 | 0.7681 | 0.8302 | 1.0000 | 0.7887 | 0.8113 | 0.8571 | 0.7765 | 0.8679 | 1.0000 |
| 66 | T0993s1-D1 | 0.8500 | 0.8500 | 0.9808 | 0.7500 | 0.7500 | 0.9231 | 0.8435 | 0.9160 | 1.0000 |
| 67 | T0993s2-D1 | 0.6538 | 0.6538 | 0.7895 | 0.5000 | 0.5000 | 0.5789 | 0.8351 | 0.9583 | 1.0000 |
| 68 | T0995-D1 | 0.7640 | 0.7640 | 0.9138 | 0.6378 | 0.6378 | 0.8448 | 0.7816 | 0.9452 | 0.9828 |
| 69 | T0996-D1 | 0.8065 | 0.8065 | 1.0000 | 0.7708 | 0.7708 | 1.0000 | 0.8019 | 0.9245 | 1.0000 |
| 70 | T0996-D2 | 0.7368 | 0.7368 | 0.9600 | 0.7556 | 0.7556 | 0.9600 | 0.7603 | 0.8889 | 1.0000 |
| 71 | T0996-D3 | 0.7667 | 0.7667 | 0.8421 | 0.8800 | 0.8980 | 1.0000 | 0.7576 | 0.8776 | 0.9474 |
| 72 | T0996-D4 | 0.7955 | 0.7955 | 0.9615 | 0.6721 | 0.6721 | 1.0000 | 0.8425 | 0.9394 | 1.0000 |
| 73 | T0996-D5 | 0.6667 | 0.6667 | 0.9583 | 0.7255 | 0.7255 | 0.9583 | 0.6636 | 0.7667 | 0.9167 |
| 74 | T0996-D6 | 0.8649 | 0.8649 | 0.9000 | 0.8269 | 0.8431 | 1.0000 | 0.8058 | 1.0000 | 1.0000 |
| 75 | T0996-D7 | 0.7381 | 0.7381 | 0.9259 | 0.7692 | 0.7692 | 0.9630 | 0.7630 | 0.8986 | 1.0000 |
| 76 | T0997-D1 | 0.7000 | 0.7000 | 0.8611 | 0.8242 | 0.8242 | 0.9167 | 0.7297 | 0.8804 | 0.9444 |
| 77 | T0998-D1 | 0.5385 | 0.5385 | 0.5385 | 0.2000 | 0.2000 | 0.2000 | 0.0000 | 0.0000 | 0.0000 |
| 78 | T0999-D1 | 0.7158 | 0.7158 | 0.7662 | 0.6441 | 0.6441 | 0.7532 | 0.7896 | 0.9063 | 0.9870 |
| 79 | T0999-D2 | 0.8045 | 0.8045 | 0.9444 | 0.7407 | 0.7407 | 0.9111 | 0.8009 | 0.8982 | 0.9333 |
| 80 | T0999-D3 | 0.7250 | 0.7250 | 0.7714 | 0.4000 | 0.4000 | 0.4000 | 0.7989 | 0.9438 | 0.9714 |
| 81 | T0999-D4 | 0.7551 | 0.7551 | 0.7708 | 0.6364 | 0.6364 | 0.8542 | 0.7942 | 0.9008 | 0.9792 |
| 82 | T0999-D5 | 0.6623 | 0.6623 | 0.8070 | 0.7500 | 0.7692 | 0.9825 | 0.8362 | 0.9301 | 1.0000 |
| 83 | T1000-D0 | 0.5846 | 0.5846 | 0.6765 | 0.7294 | 0.7294 | 0.9216 | 0.6277 | 0.8359 | 0.9412 |
| 84 | T1000-D1 | 0.7895 | 0.7895 | 1.0000 | 0.8077 | 0.8780 | 1.0000 | 0.8873 | 0.9756 | 1.0000 |
| 85 | T1000-D2 | 0.4468 | 0.4468 | 0.4884 | 0.6757 | 0.6757 | 0.8140 | 0.6326 | 0.8233 | 0.9302 |
| 86 | T1001-D1 | 0.6667 | 0.6667 | 0.6667 | 0.7083 | 0.7083 | 0.7083 | 0.2500 | 0.2500 | 0.2500 |
| 87 | T1002-D0 | 0.8872 | 0.8872 | 1.0000 | 0.8788 | 0.8788 | 0.9811 | 0.7138 | 0.8582 | 0.9623 |
| 88 | T1002-D1 | 0.9167 | 1.0000 | 1.0000 | 0.9048 | 1.0000 | 1.0000 | 0.8298 | 0.9655 | 1.0000 |
| 89 | T1002-D2 | 0.9459 | 1.0000 | 1.0000 | 0.8837 | 0.9310 | 1.0000 | 0.8667 | 0.9310 | 1.0000 |
| 90 | T1002-D3 | 0.8772 | 0.8772 | 0.9643 | 0.8519 | 0.8519 | 1.0000 | 0.8671 | 0.9155 | 1.0000 |
| 91 | T1003-D1 | 0.7611 | 0.7611 | 0.8851 | 0.6377 | 0.6377 | 0.7816 | 0.7821 | 0.9450 | 0.9885 |
| 92 | T1004-D1 | 0.8800 | 0.8800 | 0.9412 | 0.8451 | 0.9286 | 1.0000 | 0.6429 | 0.6429 | 0.9412 |
| 93 | T1004-D2 | 0.8000 | 0.8000 | 0.9333 | 0.6667 | 0.6667 | 0.6667 | 0.5000 | 0.5000 | 0.5000 |
| 94 | T1004-D3 | 0.7353 | 0.7353 | 0.8222 | 0.7500 | 0.7500 | 0.8667 | 0.3448 | 0.3448 | 0.3448 |
| 95 | T1005-D1 | 0.5800 | 0.5800 | 0.5800 | 0.5303 | 0.5303 | 0.5231 | 0.6646 | 0.8333 | 0.9538 |
| 96 | T1006-D1 | 0.6923 | 0.6923 | 0.9333 | 0.9149 | 0.9474 | 1.0000 | 0.8947 | 1.0000 | 1.0000 |
| 97 | T1010-D1 | 0.5161 | 0.5161 | 0.5161 | 0.6829 | 0.6829 | 0.6829 | 0.5179 | 0.5179 | 0.6098 |
| 98 | T1011-D1 | 0.5652 | 0.5652 | 0.5652 | 0.5417 | 0.5417 | 0.5417 | 0.7035 | 0.7667 | 0.9333 |
| 99 | T1011-D2 | 0.8333 | 0.8333 | 0.9677 | 0.6863 | 0.6863 | 0.7419 | 0.5686 | 0.7975 | 0.9677 |
| 100 | T1013-D1 | 0.7742 | 0.7742 | 0.7742 | 0.6486 | 0.6486 | 0.6486 | 0.6188 | 0.7413 | 0.8947 |
| 101 | T1014-D1 | 0.6667 | 0.6667 | 0.9355 | 0.7778 | 0.7778 | 0.9677 | 0.8797 | 0.9747 | 1.0000 |
| 102 | T1014-D2 | 0.8000 | 0.8000 | 0.8000 | 0.5870 | 0.7241 | 0.7826 | 0.6018 | 0.6724 | 0.9130 |
| 103 | T1015s1-D1 | 0.8000 | 0.8000 | 0.9412 | 0.5862 | 0.5862 | 0.8235 | 0.8125 | 0.8140 | 0.9412 |
| 104 | T1015s2-D1 | 0.7805 | 0.7805 | 0.8800 | 0.7885 | 0.7885 | 1.0000 | 0.7403 | 0.7656 | 0.9200 |
| 105 | T1016-D1 | 0.7143 | 0.7143 | 0.8000 | 0.7444 | 0.7444 | 0.9250 | 0.8657 | 0.9600 | 1.0000 |
| 106 | T1017s1-D1 | 0.6000 | 0.6000 | 0.8095 | 0.6579 | 0.6579 | 0.8095 | 0.6232 | 0.6852 | 0.9524 |
| 107 | T1017s2-D1 | 0.6585 | 0.6585 | 0.7600 | 0.8772 | 0.8772 | 0.9200 | 0.5172 | 0.5172 | 0.6000 |
| 108 | T1018-D1 | 0.7342 | 0.7342 | 0.7576 | 0.7317 | 0.7317 | 0.8636 | 0.8559 | 0.9759 | 1.0000 |

